# Native cyclase-associated protein and actin from *Xenopus laevis* oocytes form a 4:4 complex with a tripartite structure

**DOI:** 10.1101/2020.10.02.323899

**Authors:** Noriyuki Kodera, Hiroshi Abe, Shoichiro Ono

## Abstract

Cyclase-associated protein (CAP) is a conserved actin-binding protein that regulates multiple aspects of actin filament dynamics, including polymerization, depolymerization, filament severing, and nucleotide exchange. Intriguingly, CAP has been isolated from different cells and tissues as an equimolar complex with actin, and previous studies have shown that a CAP-actin complex contains six molecules each of CAP and actin. Here, we successfully isolated a complex of *Xenopus* cyclase-associated protein 1 (XCAP1) and actin from oocyte extracts and demonstrated that the complex contained four molecules each of XCAP1 and actin. The XCAP1-actin complex remained stable as a single population of 340 kDa in hydrodynamic analysis using gel filtration or analytical ultracentrifugation. Examination of the XCAP1-actin complex by high-speed atomic force microscopy revealed a tripartite structure: a middle globular domain and two globular arms. The two arms were connected with the middle globular domain by a flexible linker and observed in two states with different heights, presumably representing the presence or absence of G-actin. We hypothesize that the middle globular domain corresponds to a tetramer of the N-terminal helical-folded domain of XCAP1, and that each arm in the high state corresponds to a hetero-tetramer containing a dimer of the C-terminal CARP domain of XCAP1 and two G-actin molecules. This novel configuration of a CAP-actin complex may represent a functionally important aspect of this complex.

## Introduction

Regulated assembly and disassembly of actin filaments are vital to the diverse function of the actin cytoskeleton (1). Cyclase-associated protein (CAP) is one of the actin-regulatory proteins that control multiple key aspects of actin filament dynamics (2,3). CAP was originally identified in yeast as a protein that binds to adenylyl cyclase and is involved in the Ras signaling pathway (4,5). However, CAP was later recognized as an actin-binding protein in a variety of eukaryotes. CAP binds to actin monomers and inhibits polymerization (6). CAP also promotes exchange of actin-bound nucleotides in competition with cofilin and increases ATP-bound actin monomers that are readily available for polymerization (7-9). In addition, CAP and cofilin interact with actin filaments to enhance severing (10,11) and monomer dissociation from the pointed ends (12,13). A combination of CAP and twinfilin also enhances actin monomer dissociation from filament ends (14). CAP is involved in a number of cellular events that require actin remodeling in various cell types and tissues. For example, CAP is essential for muscle sarcomere organization in *Caenorhabditis elegans* (15) and mice (16), and deficiency of CAP2, a mammalian CAP isoform, causes cardiomyopathy in mice (17,18) and humans (19).

Intriguingly, when non-recombinant native CAP is isolated from tissues or cells, actin is associated with CAP in a multimeric complex at an equimolar ratio and cannot be dissociated without harsh conditions.

Porcine CAP (originally reported as ASP-56) was isolated from platelets as a complex with actin, and actin was finally dissociated from CAP by 3M urea (20). Similar CAP-actin complex has been isolated from yeast (8), bovine thymus (11), and mouse brain (21). The CAP-actin complex promotes actin filament disassembly in the presence of cofilin (8,11). In addition, recent studies have shown that the CAP-actin complex containing acetylated actin is an inhibitor of inverted formin 2 (INF2) (21,22). Thus, the CAP and actin have biological functions as a complex, but how the complex is assembled and why the complex formation is important for its functions remain unknown.

The native complex of yeast CAP (also known as Srv2) and actin is a 6:6 complex of ∼600 kDa (8), which can be reconstituted from purified components (23). The CAP-actin complex from mouse brain is also in a similar size (21). The N-terminal half of yeast and mouse CAPs form a hexameric “*shuriken*” structure, which is mediated by oligomerization of a putative coiled-coil region at the most N-terminus (10,24) and dimerization of the helical-folded domain (HFD) (12,25,26). The C-terminal half of CAP contains a CAP and X-linked retinitis pigmentosa 2 protein (CARP) domain that dimerizes through the most C-terminal dimerization motif (27,28). The CARP domain of CAP binds to actin monomer (6,29-31), and a CARP dimer and two actin molecules form a compact globular structure (32). Although we know structures of parts of the CAP-actin complex, we still have limited knowledge on the structure of the entire complex. Furthermore, a recent study has demonstrated that the N-terminal regions of human CAP1 and CAP2 primarily form tetramers instead of hexamers (33).

Therefore, whether the 6:6 configuration is conserved among CAP-actin complexes from different sources remains unknown. In this study, we purified a complex of *Xenopus* CAP1 and actin and demonstrated that the complex contained the two proteins in a 4:4 stoichiometric ratio, which is a novel configuration of the CAP-actin complex.

## Results

### Xenopus CAP1 (XCAP1) and actin form a 4:4 complex

We purified a native complex of CAP and actin from *Xenopus* oocyte extracts (Fig. 1). When *Xenopus* oocyte extracts were applied to a column in which glutathione S-transferase (GST)-fused *Xenopus* ADF/cofilin (XAC) was immobilized, several proteins specifically bound to the column as described previously (34) (Fig. 1A). We reported that the 65-kDa, 42-kDa, and 19-kDa proteins were *Xenopus* actin-interacting protein 1 (XAIP1), actin, and XAC, respectively (34). Peptide sequencing identified that the 94-kDa and 60-kDa proteins were gelsolin (35) and cyclase-associated protein 1 (XCAP1) (36), respectively. We attempted to isolate XCAP1 using anion-exchange chromatography followed by hydroxyapatite chromatography, but XCAP1 and actin were not separated during these procedures, and were instead purified together in an equimolar ratio (Fig. 1B). Further gel filtration chromatography using Sephadex G-200 also resulted in co-elution of XCAP1 and actin in a single peak at ∼390 kDa (our unpublished observation), which is much larger than XCAP1 or actin alone, or a 1:1 complex, indicating that they form a stable multimeric complex.

**Figure 1.**
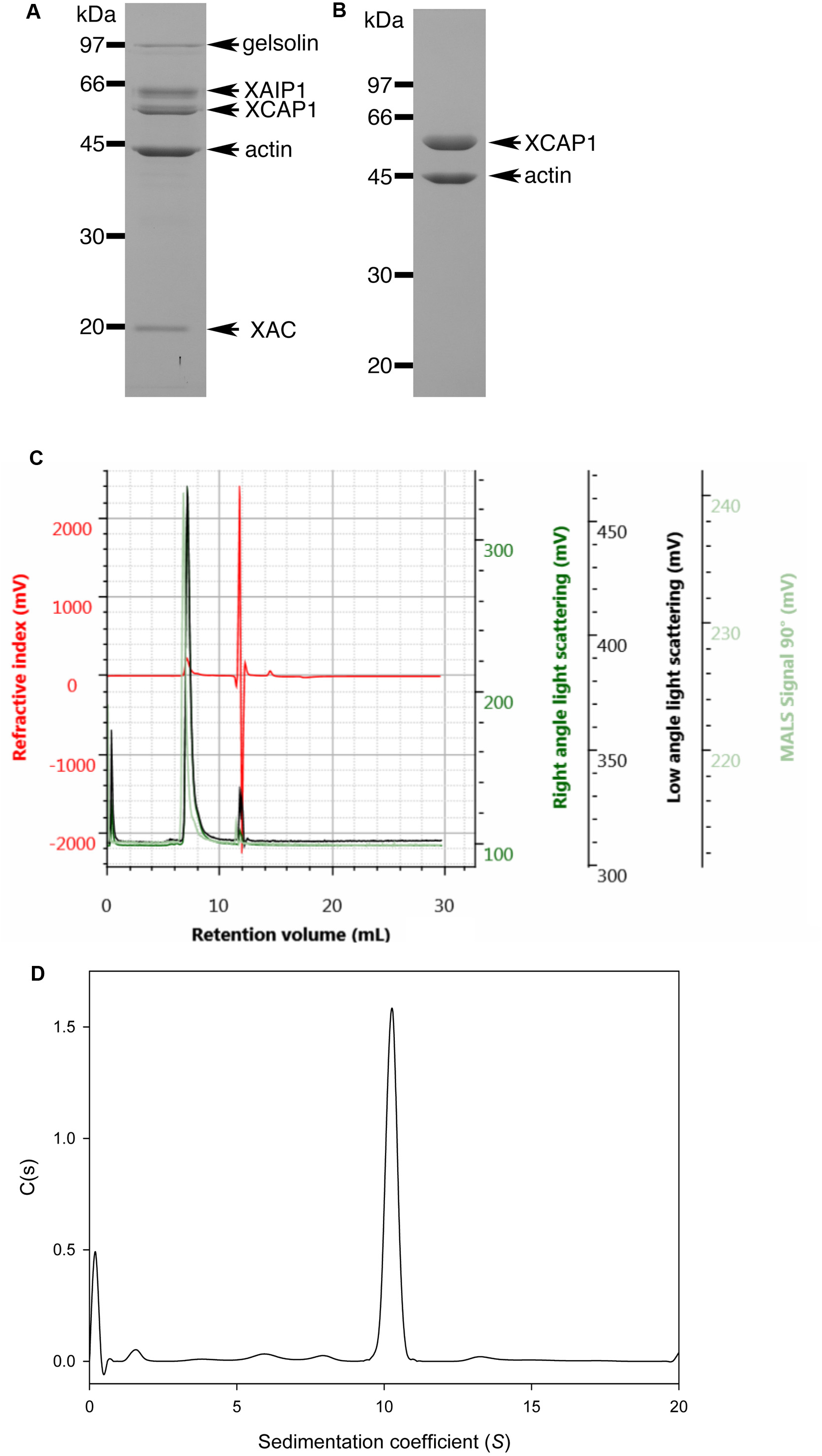
Determination of native molecular weight of the XCAP1-actin complex. (A, B) Purification of XCAP1-actin complex from *Xenopus* oocyte extracts. (A) Proteins that bound to the XAC-affinity column were eluted, separated by SDS-PAGE, and stained with Coomassie Brilliant Blue. Each band was identified by peptide sequencing as shown on the right of the gel. (B) A complex of XCAP1 and actin was isolated after anion exchange chromatography and hydroxyapatite chromatography. Positions of molecular mass markers in kDa are shown on the left. (C) SEC-MALS analysis of the XCAP1-actin complex. Purified XCAP1-actin complex was applied to size-exclusion chromatography, and refractive index (mV, red), right angle light scattering (mV, dark green), low angle light scattering (mV, black), MALS signal at 90° (mV, light green) were monitored. (D) Analytical ultracentrifugation analysis of the XCAP1-actin complex. A single peak of 10S was detected indicating that the XCAP1-actin complex was stable.

Native molecular mass of the XCAP1-actin complex was determined more accurately by two different methods: size-exclusion chromatography coupled with multi-angle light scattering (SEC-MALS) and analytical ultracentrifugation. In SEC-MALS, the XCAP1-actin complex was resolved as a single peak with a molecular mass of 340 kDa (Fig. 1C). There were no detectable peaks that corresponded to dissociated XCAP1 or actin, indicating that the XCAP1-actin complex was stable during the SEC-MALS analysis. Likewise, in analytical ultracentrifugation, the XCAP1-actin complex was resolved as a single peak of 337 kDa (*S* = 10) (Fig. 1D), which agrees with the result of SEC-MALS. Considering the molecular masses of individual XCAP1 (52 kDa) and actin (42 kDa), the native molecular mass of the XCAP1-actin complex most closely matched with that of a 4:4 complex (calculated molecular mass of 376 kDa). Since CAPs are known to bind to actin monomers, the XCAP1-actin complex most likely contains G-actin. Therefore, these results indicate that the native XCAP1 and G-actin form a stable complex at a 4:4 stoichiometric ratio.

### XCAP1-actin complex has a tripartite structure as revealed by high-speed atomic force microscopy

Structure of the XCAP1-actin complex in its native state was examined by high-speed atomic force microscopy (Fig. 2 and 3). Typical images on mica surfaces showed that the complex consisted of three globular domains (Fig. 2A), which we designated as the middle globular domain (MGD, shown in red in Fig. 2A cartoons) and two arms (Arm 1 and Arm 2, shown in green or blue in Fig. 2A cartoons). The height of MGD was 3.6 ± 0.9 nm (n = 107) (Fig. 2B, C) and remained relatively stable during time-lapse imaging (Fig. 2A and G, Supplementary Movie S1). By contrast, the two arms were observed in two different states: a high state (Arm-HS, shown in blue in Fig. 2A cartoons) and a low state (Arm-LS, shown in green in Fig. 2A cartoons, also see Fig. 3). The height of Arm-HS was 7.5 ± 0.5 nm (n = 855) (Fig. 2D and E), while that of Arm-LS was 3.3 ± 0.3 nm (n = 1078) (Fig. 2F and Fig. 3C and D). In some cases, the arms transitioned either from Arm-LS to Arm-HS (Fig. 2A and G, blue line at ∼0.9 s) or from Arm-HS to Arm-LS (Fig. 2A and G, green line at 4.5 s), suggesting that association or dissociation of a component, presumably G-actin, occurred during observations. Over the periods of HS-AFM observations, Arm-HS gradually decreased, while Arm-LS predominated, suggesting that Arm-HS was converted to Arm-LS by dissociation of actin over time likely due to repeated tapping by the AFM probe and adsorption of the complex on the surface (see below). Some of the complexes had Arm-LS throughout the observations (Fig. 2A, indicated by dashed line, Fig. 3A, Supplementary Movie S2), and the height of Arm-LS was 3.3 ± 0.3 nm (n = 1078) (Fig. 3B-D).

**Figure 2.**
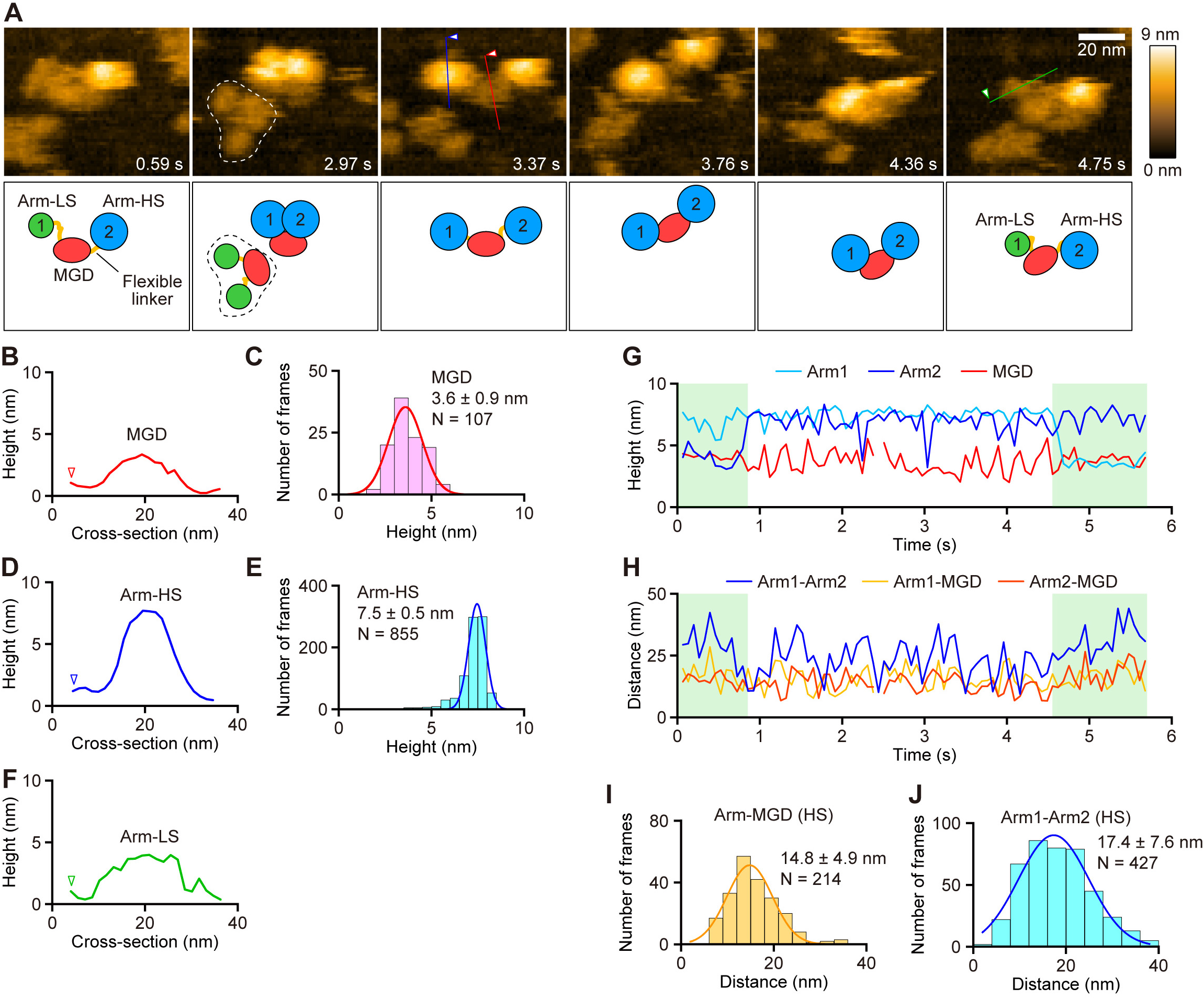
High-speed atomic force microscopy reveals a tripartite structure of the XCAP1-actin complex. (A) Time-lapse HS-AFM images of the XCAP1-actin complex on a mica surface (see Supplementary Movie S1). Scanning area was 80 x 64 nm^2^ with 64 x 48 pixels. Imaging rate was 66 ms/frame (∼15 fps). Bar, 20 nm. Schematic representation of molecular features is shown in the bottom panels: middle globular domain (MGD, red), arm in the low state (Arm-LS, green), and arm in the high state (Arm-HS, blue). The complex indicated by dashed lines in the second frame had both arms in Arm-LS throughout the observation (see Fig. 3 for quantitative analysis). (B-F) Cross-section analyses of MGD (B, red), Arm-HS (D, blue), and Arm-LS (F, green) at the straight colored lines drawn on the images in A. Height distributions of MGD (C) and Arm-HS and single Gaussian fitting yielded average heights of MGD and Arm-HS as indicated in the figure. (G) Time course of the heights of three globular domains. Green-shaded areas indicate periods when one of the arms were in the low state. (H-J) Time course of the distances between the domains at their highest points (H). Distribution of the distance between two arms (I), and between MGD and one of the arms (J) and single Gaussian fitting yielded average distances as indicated in the figure. Arm-HS was selected in these analyses.

**Figure 3.**
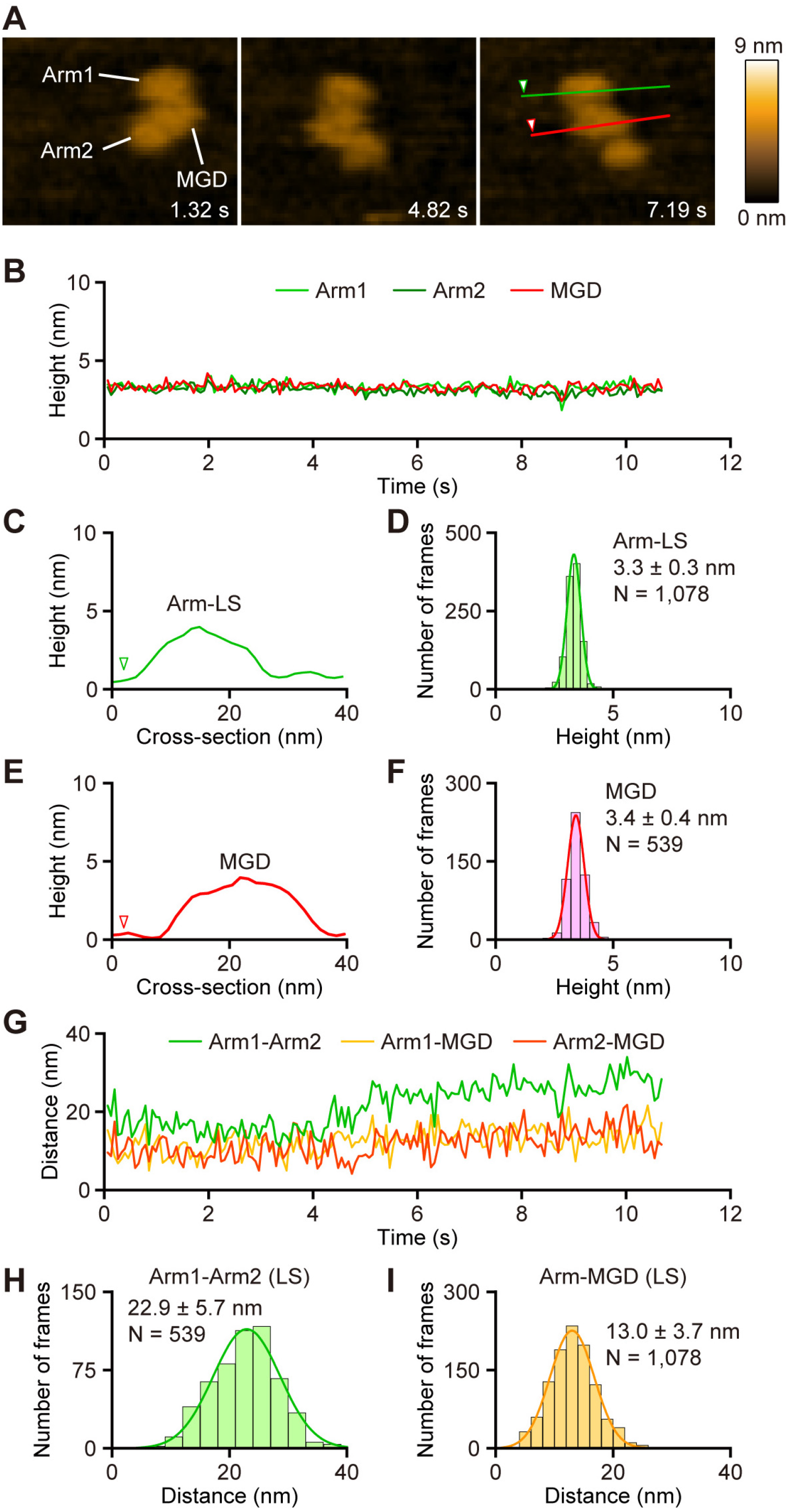
The XCAP1-actin complex with both arms in a low state is stable. (A) Time-lapse HS-AFM images of the XCAP1-actin complex containing both arms in Arm-LS on a mica surface (see Supplementary Movie S2). Scanning area was 80 x 64 nm^2^ with 64 x 48 pixels. Imaging rate was 66 ms/frame (∼15 fps). Bar, 20 nm. (B) Time course of the heights of three globular domains. (C-F) Cross-section analyses of Arm-LS (C, green) and MGD (E, red) at the straight colored lines drawn on the image in A. Height distributions of Arm-LS (D) and MGD (F) and single Gaussian fitting yielded average heights of Arm-LS and MGD as indicated in the figure. (G-I) Time course of the distances between the domains at their highest points (G). Distribution of the distance between two arms (H), and between MGD and one of the arms (I) and single Gaussian fitting yielded average distances as indicated in the figure.

The two arms were very dynamic when they were in Arm-HS (Fig. 2A. Supplementary Movie S1), but restricted within 14.8 ± 4.9 nm (n = 214) of MGD (Fig. 2H and I) as if the arms were connected to MGD by flexible linkers. The distance between the highest points of the two arms in Arm-HS fluctuated in a wide range with an average of 17.4 ± 7.6 nm (n = 427) (Fig. 2H and J), further supporting the presence of flexible linkers between MGD and each arm. However, once arms were converted from Arm-HS to Arm-LS, they were stabilized in Arm-LS (Fig. 3A-D, Supplementary Movie S2), while MGD remained unchanged (Fig. 3B, E, and F). The distance between two arms became wider [22.9 ± 5.7 nm (n = 539)], whereas that between each arm and MGD became narrower [13.0 ± 3.7 nm (n = 1078)]. These results suggest that Arm-LS was physically stabilized by adsorption to the mica surface.

To test how surface adsorption affects the molecular features of the XCAP1-actin complex, we used a mica surface that was treated with APTES, which adds positive charges to the surface and causes non-specific strong adsorption of proteins. On APTES-treated mica, the two arms were almost always detected in the low state (Arm-LS) with the height of 3.0 ± 0.4 nm (n = 2293) (Fig. 4A-D, Supplementary Movie S3). The height and shape of MGD were indistinguishable between normal and APTES-treated mica surfaces (Fig. 4A, B, E, and F). The distance between the two arms remained relatively constant at 25.4 ± 5.6 nm (n = 1331) (Fig. 4G and H), which is much wider than that of two arms in Arm-HS on normal mica surfaces (Fig. 2I), suggesting that the two arms were strongly immobilized on the surface and spread apart. By contrast, the distance between MGD and an arm remained constant on the APTES-treated surface (Fig. 4G and I) in a similar manner to MGD and Arm-LS on the normal surface (Fig. 3I). These observations suggest that strong adsorption of the XCAP1-actin complex onto a solid surface artificially converts Arm-HS to Arm-LS by causing dissociation of an arm-bound component, which we hypothesize to be G-actin.

**Figure 4.**
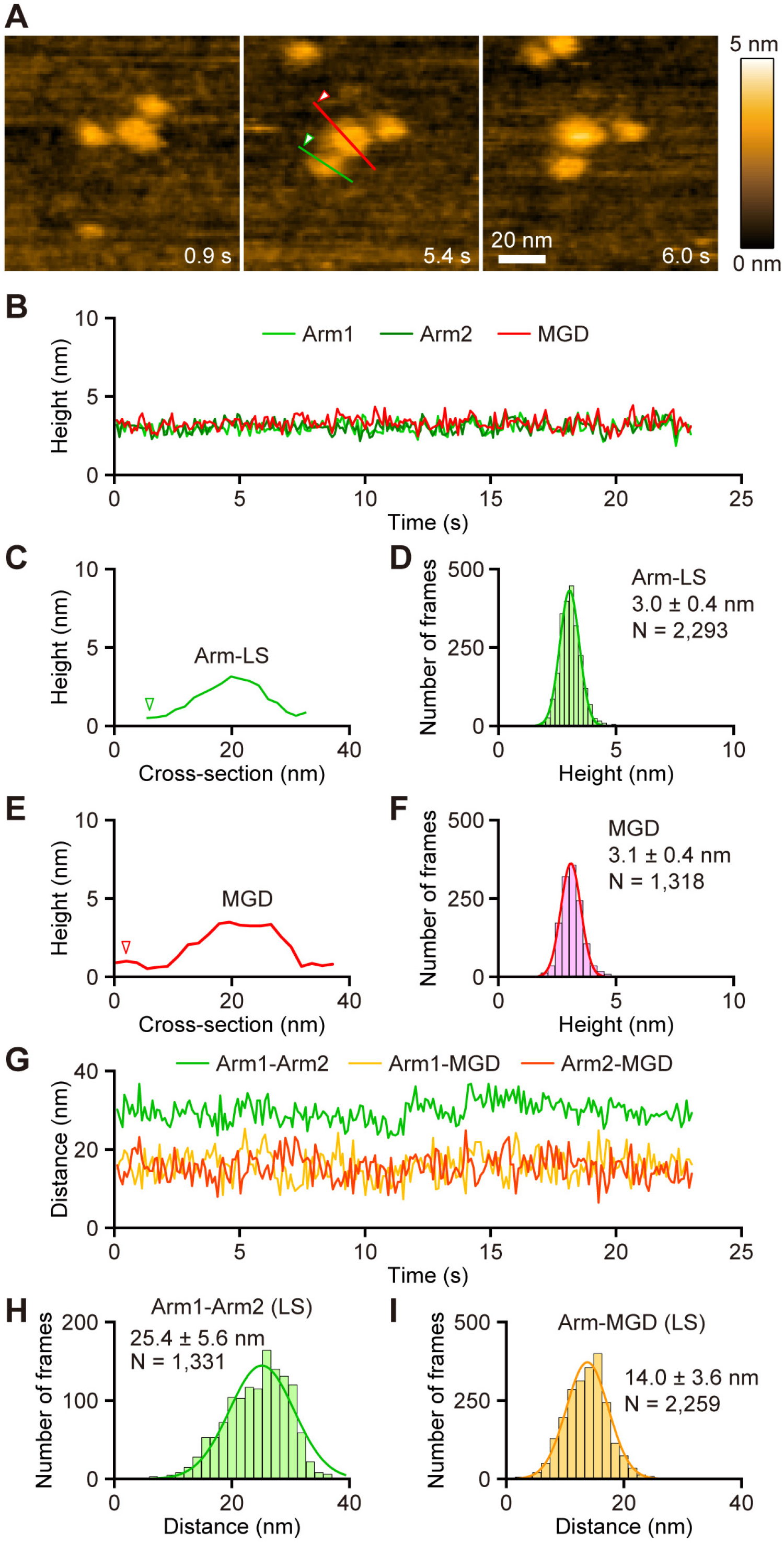
Strong adsorption of the XCAP1-actin complex to a charged surface stabilizes two arms in a low state. Time-lapse HS-AFM images of the XCAP1-actin complex on an APTES-treated mica surface (see Supplementary Movie S3). Scanning area was 100 ⨯ 100 nm^2^ with 80 x 80 pixels. Imaging rate was 100 ms/frame (10 fps). Bar, 20 nm. (B) Time course of the heights of three globular domains. (C-F) Cross-section analyses of Arm-LS (C, red) and Arm-LS (E, green) at the straight colored lines drawn on the image in A. Height distributions of Arm-LS (D) and MGD (F) and single Gaussian fitting yielded average heights of MGD and Arm-HS as indicated in the figure. (G-I) Time course of the distances between the domains at their highest points (G). Distribution of the distance between two arms (H), and between MGD and one of the arms (I) and single Gaussian fitting yielded average distances as indicated in the figure.

## Discussion

Based on known biochemical and biophysical properties of CAP from other species, we propose a model for the structure of the XCAP1-actin complex, which is in the appearance of two “butterflies” (CARP/G-actin) and a “flower” (HFD) (Fig. 5). We hypothesize that MGD corresponds to a tetramer of the HFD of XCAP1, and that each arm domain in the high state (Arm-HS) corresponds to a hetero-tetramer containing a dimer of the CARP domain of XCAP1 and two G-actin molecules (Fig. 5). The HFD of CAP by itself forms a dimer (12,25,26), and the N-terminal oligomerization motif forms a putative coiled-coil and mediates formation of a tetramer (33) or hexamer (10,23,24) of the HFD. The height of MGD (Fig. 2C, 3F, 4F) matches with that of the diameter of one HFD (12,25), suggesting that each HFD is laterally attached to the substrate. By contrast, the C-terminal dimerization motif mediates rigid dimerization of the CARP domain through strand-exchange β-sheet formation (27), which then binds to two G-actin molecules (32). Again, the height of Arm-HS (Fig. 2E) matches with the diameter of the hetero-tetramer of CARP and G-actin (32). Proline-rich regions and Wiskott Aldrich Syndrome protein-homology 2 (WH2) can serve as a flexible linker between HFD and CARP (Fig. 5). WH2 of CAP binds to G-actin (31,37), but transient dissociation of WH2 from G-actin can allow full extension of the linker with structural flexibility. WH2 of CAP also binds to the N-terminal diaphanous inhibitory domain of INF2 (22). Therefore, flexibility of WH2 in the CAP-actin complex should keep it accessible with INF2. This hypothetical architecture of the XCAP1-actin complex needs to be tested by additional structural analysis at higher resolutions or localization of components using specific probes.

**Figure 5.**
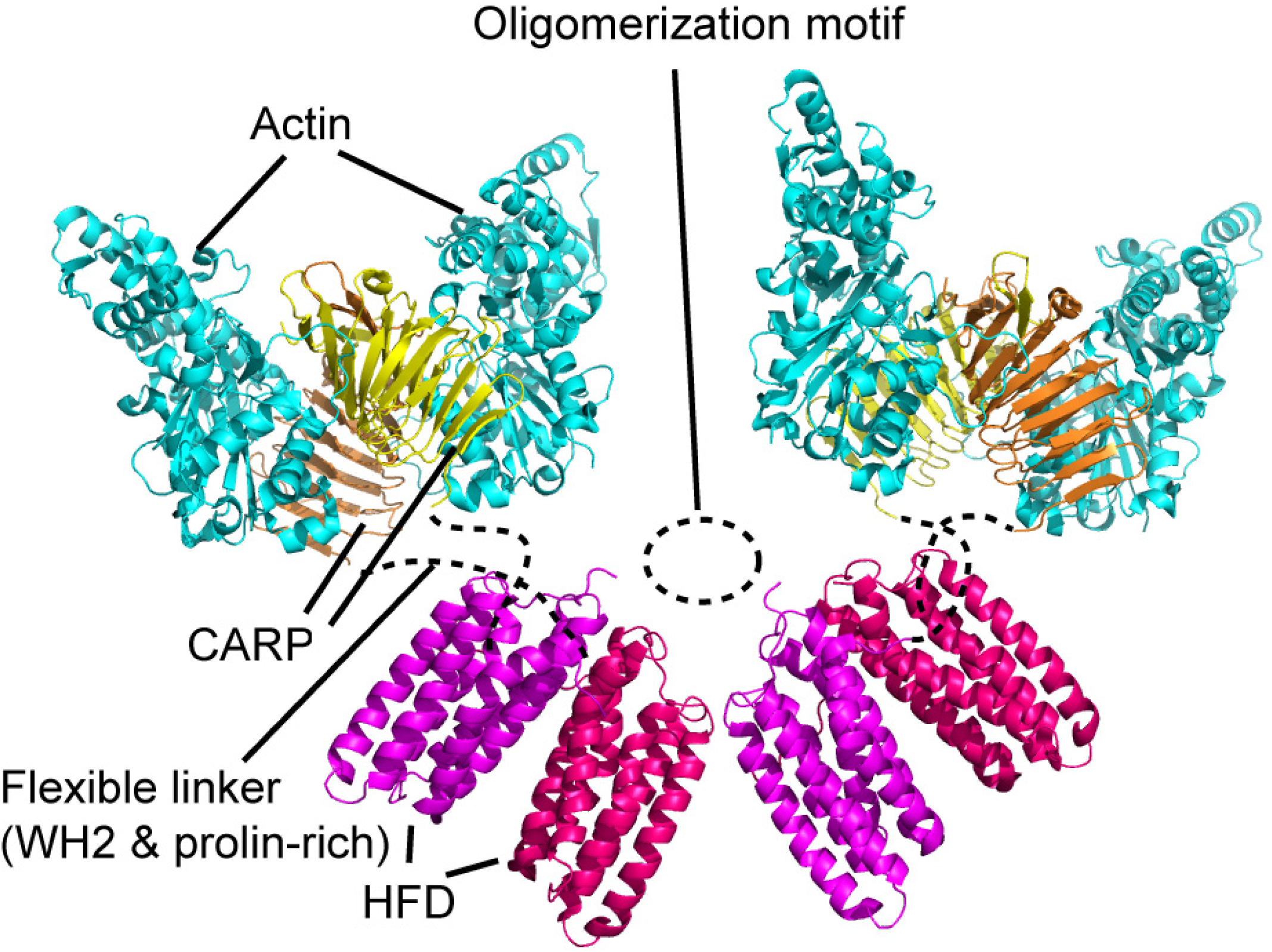
Model of the CAP-actin complex. Crystal structures of HFD of mouse CAP1 (Protein Data Bank accession ID: 6RSQ) and CARP domain of mouse CAP1 bound to actin (Protein Data Bank accession ID: 6FM2) were used to reconstruct a CAP-actin complex at a 4:4 stoichiometric ratio. Putative locations of oligomerization motif and flexible linkers are indicated by dashed lines.

The configuration of the XCAP1-actin complex in a 4:4 stoichiometric ratio is different from other reported configurations of CAP-actin complexes from different organisms in a 6:6 molar ratio (8,23). It is worth noting that the tripartite structure of the XCAP1-actin complex is very similar to one of the electron microscopy images of yeast Srv2/CAP-actin complex (see Fig. 1 of Ref. (8)). However, the yeast Srv2/CAP-actin complex is a 600-kDa complex containing Srv2/CAP and actin in a molar ratio of 6:6 (8). It remains unclear whether such a difference represents an inherent structural variety of the complex assembly or technical artifacts in the analyses or preparation methods. Further studies are needed to test whether the XCAP1-actin complex with a similar configuration can be reconstituted from purified XCAP1 and G-actin, which should allow dissection of domains and residues that are required for assembly of the XCAP1-actin complex.

Our structural model of the CAP-actin complex places the actin-binding site of HFD outward, suggesting that any two of the four HFDs can interact with two actin subunits at the pointed end of a filament to accelerate depolymerization (12,13). It allows two remaining free HFD to interact with newly exposed actin subunits at the pointed end, such that CAP can stay bound to a filament and processively depolymerize actin filaments from the pointed end.

However, whether CAP can depolymerize actin filaments processively remains to be determined. In addition, the CARP domain, which has nucleotide exchange activity, is in proximity and available to capture newly depolymerized ADP-actin and promote rapid conversion to ATP-actin. It would be interesting to determine whether binding of other proteins to WH2 or proline-rich region in the flexible linker (22,38-40) or phosphorylation of CAP (41,42) alters the structure and function of the CAP-actin complex. Thus, our structural model provides mechanistic insight in the function of CAP in the regulation of actin turnover.

## Experimental procedures

### Purification of XCAP1-actin complex from Xenopus laevis oocytes

Extracts from *Xenopus laevis* oocytes were prepared and applied to an affinity column in which GST-fused *Xenopus* ADF/cofilin (XAC) had been immobilized as described (34). Proteins bound to the column were eluted with 1 M NaCl, 2 mM MgCl_2_, 1 mM dithiothreitol (DTT), 0.01% NaN_3_, and 20 mM HEPES-KOH, pH 7.2. The eluate was fractionated with ammonium sulfate at 45% saturation. The precipitates obtained by centrifugation was dissolved and dialyzed against 60 mM NaCl buffer (60 mM NaCl, 0.5 mM DTT, 0.01% NaN_3_, and 20 mM Tris-HCl, pH 8.0), applied to a DE52 column pre-equilibrated with the same buffer, and then eluted with a linear gradient of 60-300 mM NaCl. The fractions containing XCAP1-actin complex was directly applied to a hydroxyapatite column pre-equilibrated with 60 mM NaCl buffer and washed thoroughly with the same buffer. XCAP1-actin complex was eluted with a linear gradient of 0-300 mM potassium phosphate buffer at pH 8.0. Purified XCAP1-actin complex was concentrated by ultrafiltration with Ultracel-30K (Millipore) and dialyzed against 0.1 M KCl, 2 mM MgCl_2_, 1 mM DTT, 0.01% NaN_3_, and 20 mM HEPES-KOH, pH 7.2.

### Size exclusion chromatography - multi-angle light scattering

Samples were analyzed by a Malvern OmniSEC integrated system (Malvern Pananalytical, MA) with a SRT SEC-300 (Sepax, Newark, DE) analytical SEC column. Samples were loaded from an autoinjector sample tray that was kept at 20 °C. Phosphate-buffered saline (pH 7.2) containing 0.02 % NaN_3_ was used in a mobile phase. Calibration was done using a bovine serum albumin standard. Data from a refractive index, right angle light scattering (RALS), low angle light scattering (LALS), viscosimeter and a UV PDA detector were collected. The resulting chromatograms were analyzed using triple detection (RI, RALS and viscosimeter) and the dn/dc from sample concentration were used to calculate molecular weight of the peaks as well as hydrodynamic radius. Molecular weights were calculated with Malvern OmniSEC software version 10.41.

### Analytical ultracentrifugation

Sedimentation velocity data were collected using a Beckman Optima AUC analytical ultracentrifuge using a rotor speed of 40,000 rpm (128794 g) at 20 °C. Data were recorded by monitoring the sedimentation of the absorbance at 280 nm using a radial step size of 0.001 cm. Set parameters included a partial specific volume (Vbar) of 0.73, a buffer viscosity of 1.002 Poise, and density of 1.00 g/ml. Sedimentation velocity data were analyzed using both SEDFIT (www.analyticalultracentrifugation.com) (43) and UltraScan (www.ultrascan.uthscsa.edu) (44). Continuous sedimentation coefficient distribution c(*s*) analyses were restrained by maximum entropy regularization at *P* = 0.95 confidence interval. The baseline, meniscus, frictional coefficient, systematic time-invariant and radial invariant noise were fit.

### High-speed atomic force microscopy

HS-AFM imaging was performed in solution at room temperature using a laboratory-built HS-AFM setup (45,46), as described previously (47). In brief, a glass sample stage (diameter, 2 mm; height, 2 mm) with a thin mica disc (1.5 mm in diameter and ∼0.05 mm in thickness) glued to the top by epoxy was attached onto the top of a Z-scanner by a drop of nail polish. Either bare mica surface or APTES ((3-aminopropyl)triethoxysilane) treated mica surface (48) was used as a substrate. Onto either substrate, a drop (2 μl) of diluted protein sample (ca. 1 nM) with buffer A (100 mM KCl, 20 mM HEPES–KOH, pH 7.2, 2 mM MgCl_2_) was deposited for 3 min. For the observation using bare mica surface, the surface was rinsed with 20 μl of buffer B (30 mM KCl, 20 mM HEPES–NaOH, pH 7.0, 2 mM MgCl_2_) and imaged in buffer B. For the observation using APTES treated mica surface, the surface was rinsed with 20 μl of buffer A and imaged in buffer A. AFM imaging was carried out in a tapping mode using small cantilevers (BLAC10DS-A2, Olympus) (resonant frequency, ∼0.5 MHz in water; quality factor, ∼1.5 in water; Spring constant, ∼0.1 N m^−1^). The probe tip was grown on the original tip end of a cantilever through electron beam deposition using ferrocene and was further sharpened using a radio frequency plasma etcher (Tergeo, PIE Scientific LLC., USA) under an argon gas atmosphere (Direct mode, 10 sccm and 20 W for 1.5 min). The cantilever’s free oscillation amplitude *A*_0_ and set-point amplitude *A*_s_ were set at ∼2 nm and ∼0.9 × *A*_0_, respectively. The imaging rate, scan size, and pixel size for each AFM image are described in the figure legends.

### Data analyses of HS-AFM images

HS-AFM images were viewed and analyzed using the laboratory built software, Kodec4.4.7.39 (49). In brief, a low-pass filter to remove spike noise and a flattening filter to make the xy-plane flat were applied to individual images. The position and height of the peak within each domain were determined semi-automatically using the following steps. First, the most probable highest point was selected manually.

Second, the actual highest point was automatically determined by searching a 5 × 5 pixel area (typically 6.25 × 6.25 nm^2^) around the selected point.

### Molecular graphics

A model of the CAP-actin complex was generated using PyMol (Schrödinger, LLC) and annotated using Adobe Illustrator CS2 (Adobe).

## Data availability

All data are contained in the manuscript. Raw data are available from S. O. upon request.

### Acknowledgements

We thank Dr. Toshio Ando for support in HS-AFM. We wish to acknowledge the core facilities at the Parker H. Petit Institute for Bioengineering and Bioscience at the Georgia Institute of Technology for the use of their shared equipment, services and expertise. We thank Dr. Bettina Bommarius for performing SEC-MALS and AUC analyses at the Biopolymer Characterization Core.

## Abbreviations

APTES: (3-aminopropyl)triethoxysilane
Arm-HS: arm in a high state
Arm-LS: arm in a low state
CAP: cyclase-associated protein
CARP: CAP and X-linked retinitis pigmentosa 2 protein
HS-AFM: high-speed atomic force microscopy
HFD: helical folded domain
INF2: inverted formin 2
MGD: middle globular domain
SEC-MALS: size exclusion chromatography-multi angle light scattering
WH2: Wiskott Aldrich Syndrome Protein homology 2
XAC: *Xenopus* actin depolymerizing factor/cofilin
XCAP1: *Xenopus* cyclase-associated protein 1

